# Strong paracrine effects of SASP from senescence-induced severe early-onset COPD-derived fibroblasts

**DOI:** 10.1101/2023.09.01.555721

**Authors:** R.A. Meuleman, W. Timens, W. Kooistra, M. van den Berge, C-A. Brandsma, R.R. Woldhuis

## Abstract

**Background:** Lung fibroblasts from Severe Early-Onset (SEO-)COPD patients exhibit increased cellular senescence with higher levels of senescence-associated secretory phenotype (SASP) protein secretion. Yet, the impact of senescent fibroblasts, and their SASP, on surrounding fibroblasts in SEO-COPD lungs remains unclear.

**Aim:** To identify the effect of the SASP secreted by senescent SEO-COPD-derived fibroblasts on surrounding lung fibroblasts.

**Methods:** Cellular senescence was induced in lung fibroblasts derived from seven SEO-COPD patients (age≤53 years, FEV1<40% predicted), and conditioned medium (CM) containing the SASP, was collected (senescent CM). CM from untreated fibroblasts was used as control. Fibroblasts were stimulated with senescent and control CM, and with tissue plasminogen activator (t-PA), a previously identified COPD-associated SASP protein. Effects on paracrine senescence, inflammation, and extracellular matrix (ECM) regulation were assessed.

**Results:** Stimulation with senescent CM increased the percentage of Senescence-associated beta-galactosidase positive fibroblasts and decreased p16, p21 & *LMNB1* expression (p<0.05). T-PA did not affect these markers. Senescent CM increased IL-8 gene expression, increased IL-6 secretion, and strongly increased IL-8 secretion compared to control CM. T-PA slightly decreased IL-6 and IL-8 secretion. Additionally, senescent CM and t-PA stimulation both reduced decorin secretion. Senescent CM reduced *FN1* gene expression, while *DCN* and *MMP2* expression remained unaffected. T-PA did not affect ECM gene expression.

**Conclusion:** The SASP from senescence-induced SEO-COPD-derived fibroblasts has a strong paracrine effect on untreated fibroblasts, suggesting that senescent lung fibroblasts contribute to chronic inflammation and ECM dysregulation. These findings imply involvement of senescent fibroblasts in abnormal lung ageing and possibly disease pathology in COPD.

## Introduction

Chronic Obstructive Pulmonary Disease (COPD) is a chronic, progressive lung disease that was globally the third leading cause of death in 2019(1). Noxious gases, including cigarette smoke, contribute to the development of COPD(2, 3). Lungs of COPD patients have features of chronic inflammation, airway remodeling and loss of elastic coil(4). Yet, the exact mechanisms underlying COPD and driving disease progression remain to be elucidated.

COPD has been proposed to represent abnormal lung ageing(5). Severe early-onset (SEO-)COPD, a COPD subtype in which patients develop severe COPD at a relatively young age, is hypothesized to reflect accelerated lung ageing(6). Lung ageing is characterized by several hallmarks, including cellular senescence and extracellular matrix (ECM) dysregulation(5, 7). Several ECM alterations are observed in COPD-derived lungs, including increased levels of ECM degrading enzymes, elastic fiber breakdown and altered proteoglycan levels(8–10). Cellular senescence is an irreversible state in which cells no longer divide, but still are metabolic active(11). Previously, our group has shown increased cellular senescence in COPD-derived fibroblasts compared to fibroblasts from non-COPD ex-smokers(12). Elevated cellular senescence in these fibroblasts was associated with altered ECM production, including reduced decorin (DCN) expression and secretion(12), indicating that accelerated ageing through increased cellular senescence may contribute to ECM dysregulation in COPD lungs.

Senescent cells can exhibit detrimental effects on their environment, mainly via their secreted factors, i.e. senescence-associated secretory phenotype (SASP), which includes pro-inflammatory cytokines, growth factors, ECM degrading enzymes, serine proteases and the ECM glycoprotein fibronectin (FN1)(13). To gain insight into how senescent lung fibroblasts affect their environment and contribute to tissue dysfunction in COPD, we previously identified their SASP(14). COPD-derived fibroblasts showed higher secretion of 42 SASP proteins, compared to matched controls, including several pro-inflammatory cytokines, contributing to inflammation in COPD(14). Furthermore, this study identified tissue Plasminogen Activator (t-PA) as SASP component among the proteins with the strongest increase in SEO-COPD-derived fibroblasts compared to controls(14). T-PA is of particular interest as it can regulate levels and activity of ECM degrading enzymes, including Matrix Metalloproteinase 2 (MMP2), and facilitates fibrinolysis and thrombolysis trough activation of plasminogen(15–18). Intriguingly, the majority of the identified SASP proteins were upregulated in SEO-COPD-derived fibroblasts, suggesting that the detrimental effect of senescent fibroblasts and their SASP is especially relevant in this COPD subtype(14). Yet, it remains unknown whether fibroblasts-derived SASP could affect surrounding fibroblasts and contribute to chronic inflammation and ECM dysregulation in (SEO-)COPD lungs.

Therefore, this study aimed to investigate whether senescent fibroblasts contribute to pathological changes in COPD lungs. Since the SASP is crucial for the paracrine signaling by these cells, we aimed to study how SASP secreted by senescence-induced SEO-COPD-derived fibroblasts affects the surrounding fibroblasts. To investigate this effect, we stimulated untreated SEO-COPD-derived fibroblasts with conditioned medium (CM) from senescence-induced SEO-COPD-derived fibroblasts and compared the effects to fibroblasts stimulated with CM from untreated SEO-COPD-derived fibroblasts. Furthermore, we stimulated the fibroblasts with the SASP component t-PA. After stimulation, the effects on paracrine senescence, inflammation, and ECM regulation were assessed.

## Methods

### Full details of materials and methods are provided in the online supplement

#### Primary parenchymal lung fibroblasts

Primary lung fibroblasts were isolated as described before(19) from explant material of seven SEO-COPD patients (age ≤53 years and Forced Expiratory Volume in one second (FEV1) <40% predicted at moment of lung transplantation) who did not have alpha-1-antitrypsin deficiency(6). After isolation, fibroblasts were stored at -180°C until use. Subject characteristics are summarized in Supplementary Table S1.

#### Cell culture & senescence induction

Primary lung fibroblasts were cultured in T25-flasks (Greiner Bio-One, Frickenhausen, Germany) in Ham’s F12 medium (ThermoFisher Scientific, Waltham, USA) with 10% Fetal Calf Serum (FCS) (Lonza, Basel, Switzerland), L-glutamine (Lonza), and 1% Penicillin/Streptomycin (Lonza) (in short 10% FCS medium) from passage 3 till 5. At passage 5, 180,000 cells were seeded per T25-flask. After two days, fibroblasts were treated with 250μM paraquat (PQ) (Sigma-Aldrich, Darmstadt, Germany) or 0.2μM doxorubicin (Dox) (Selleck Chemicals, Houston, US) to induce senescence and untreated cells were used as control (Ctrl). After 24 hours, PQ and Dox were washed away, 5% FCS medium was added, and cells were cultured for four days until senescent state. Senescence-induced and untreated fibroblasts were reseeded into 12-well (Conditioned Medium (CM) collection) or 24-well plates (CM stimulation) (Corning, Corning, US) for the experiments described below.

#### Collection of conditioned medium

For CM collection, 100,000 Ctrl, 250μM PQ and 0.2μM Dox treated fibroblasts were seeded into 12-well plates to equalize cell numbers (passage 6). After 24 hours, medium was refreshed with 5% FCS medium to discard non-adherent cells and 48 hours later CM was collected. CMs were centrifuged (190g, 5 min) to discard cell debris and supernatants were directly used for CM treatment. CM leftovers were stored at -80°C.

#### Stimulation with conditioned medium

In parallel, 12,500 untreated fibroblasts were seeded into 24-well plates (passage 6) for CM stimulation. Three days after seeding, cells were stimulated with 100% CM or 50% CM (1:1 mixed with fresh 5% FCS medium) from Ctrl, 250μM PQ or 0.2μM Dox treated cells from the same donor. After 24 hours of stimulation, RNA was collected. After four days, supernatants were collected, and Senescence-associated beta-galactosidase (SA-β-gal) staining was performed.

#### Stimulation with tissue Plasminogen Activator

For tissue plasminogen activator (t-PA) stimulation, 12,500 untreated fibroblasts were seeded into 24-well plates (passage 6). Three days later, fibroblasts were treated with 0.5nM (CM concentration), 10nM or 20nM t-PA (literature-based concentrations(20–22)) (Sigma-Aldrich) or untreated as control. After 24 hours, RNA was collected. After four days, supernatants were collected, and SA-β-gal staining was performed.

### Statistical analyses

Wilcoxon matched pairs signed-rank test was performed for pairwise comparison between treatment and control samples using GraphPad Prism 8 Software. P-values <0.05 were considered significant.

## Results

### Senescent CM increases SA-β-gal and decreases p16, p21 and *LMNB1*

Conditioned medium was collected from untreated fibroblasts (Ctrl CM), and PQ-induced and Dox-induced senescent fibroblasts (Senescent CM). Senescence induction in PQ and Dox treated CM producing cells was confirmed by increased percentages of SA-β-gal positive cells (Supplementary Figure S1).

Upon CM treatment we assessed the effect on paracrine senescence using SA-β-gal staining, cell proliferation and p21, p16 and *LMNB1* gene expression. Fibroblasts treated with senescent CM showed a significant increase in SA-β-gal positive cells compared to Ctrl CM stimulated cells with both 50% and 100% of CM (Figure 1A), while cell numbers remained unaffected (Figure 1B). Both 50% and 100% PQ CM decreased p21 and p16 gene expression, and only 100% PQ CM reduced *LMNB1* expression (Figure 1C-E). Dox CM did not affect expression of these genes (Figure 1C-E). T-PA stimulation did not affect the measured senescence markers (Supplementary Figure S2).

**Figure 1.**
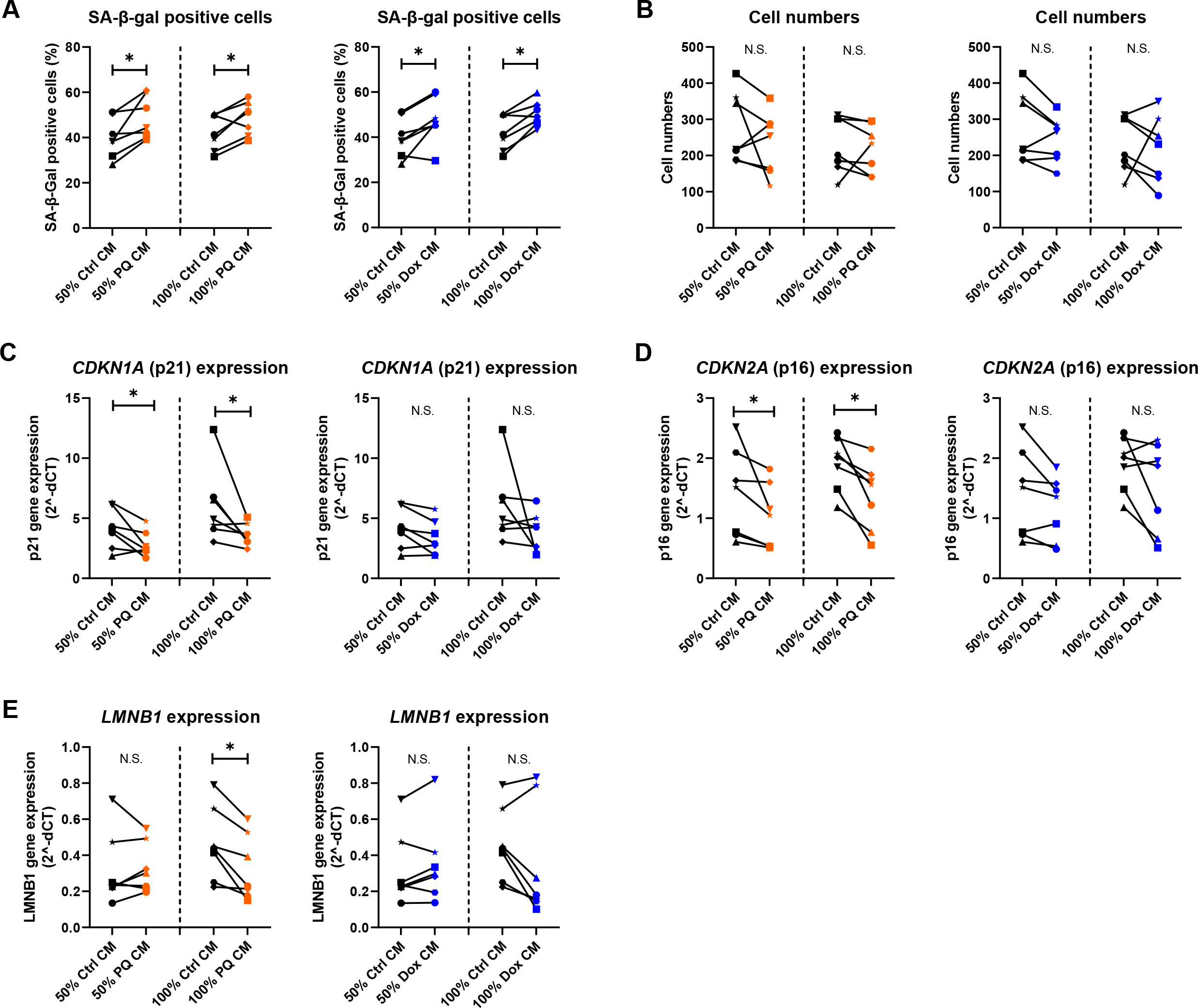
The effect of senescent CM on paracrine senescence. Untreated SEO-COPD-derived fibroblasts were stimulated with control or senescent (PQ or Dox) conditioned medium (CM). The percentage of Senescence-Associated Beta-Galactosidase (SA-β-gal) staining positive cells **(A)** and cell numbers **(B)** were assessed after four days. p21 (*CDKN1A*) **(C)**, p16 (*CDKN2A*) **(D)** and *LMNB1* **(E)** gene expression was measured after 24 hours. Wilcoxon matched pairs signed-rank test was applied for statistical testing. * p<0.05.

### Senescent CM induces a pro-inflammatory response in SEO-COPD-derived fibroblasts

Next, we assessed the effect of SASP on pro-inflammatory cytokines interleukin (IL-)6 and IL-8. After 24 hours, stimulation with senescent CM increased IL-8 gene expression, while IL-6 gene expression remained unaffected (Figure 2A & B). After four days, senescent CM stimulation increased IL-6 secretion and induced a strong increase in IL-8 secretion, which was most pronounced with 50% CM (Fold changes: 7.0 and 4.1 for 50% and 100% PQ CM resp. and 7.0 and 4.7 for 50% and 100% Dox CM resp.) (Figure 2C & D). Stimulation with 0.5nM t-PA resulted in a small decrease in IL-6 secretion only, while 10nM and 20nM t-PA only slightly decreased IL-8 secretion (Supplementary Figure S3)

**Figure 2.**
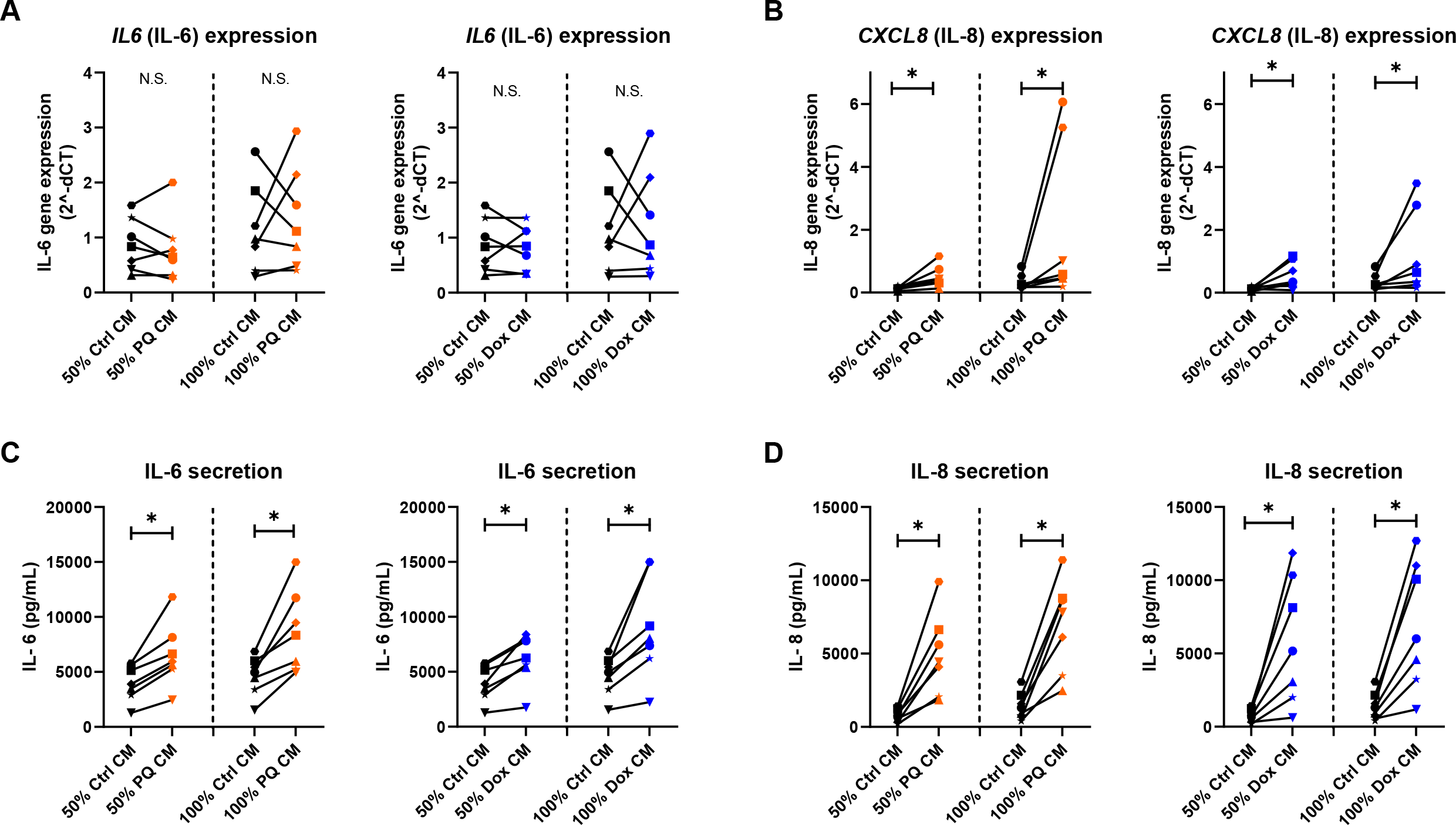
The effect of senescent CM on pro-inflammatory interleukin secretion. Untreated SEO-COPD-derived fibroblasts were stimulated with control or senescent (PQ or Dox) conditioned medium (CM). After 24 hours, IL-6 (*IL6*) **(A)** and IL-8 (*CXCL8*) **(B)** gene expression was measured. After four days, IL-6 **(C)** and IL-8 **(D)** secretion was assessed in cell culture medium using ELISA. Wilcoxon matched pairs signed-rank test was applied for statistical testing. * p<0.05.

### Senescent CM and t-PA stimulation reduced secreted DCN levels

To investigate the effect of SASP on ECM remodeling, we measured the following ECM-related genes: *DCN* as it was previously associated with COPD and senescence(12, 19), *FN1* as an early fibrotic marker(23) and *MMP2* as it can be induced by t-PA(17). *MMP2* (Supplementary Figure S4A) and *DCN* (Figure 3A) expression remained unaffected by senescent CM stimulation after 24 hours, while *FN1* gene expression was slightly decreased after stimulation with 50% PQ CM (Supplementary Figure 4B). Yet, a clear decrease in secreted DCN levels was observed after four days of senescent CM stimulation, independent of CM concentration or senescence inducer (Figure 3B).

**Figure 3.**
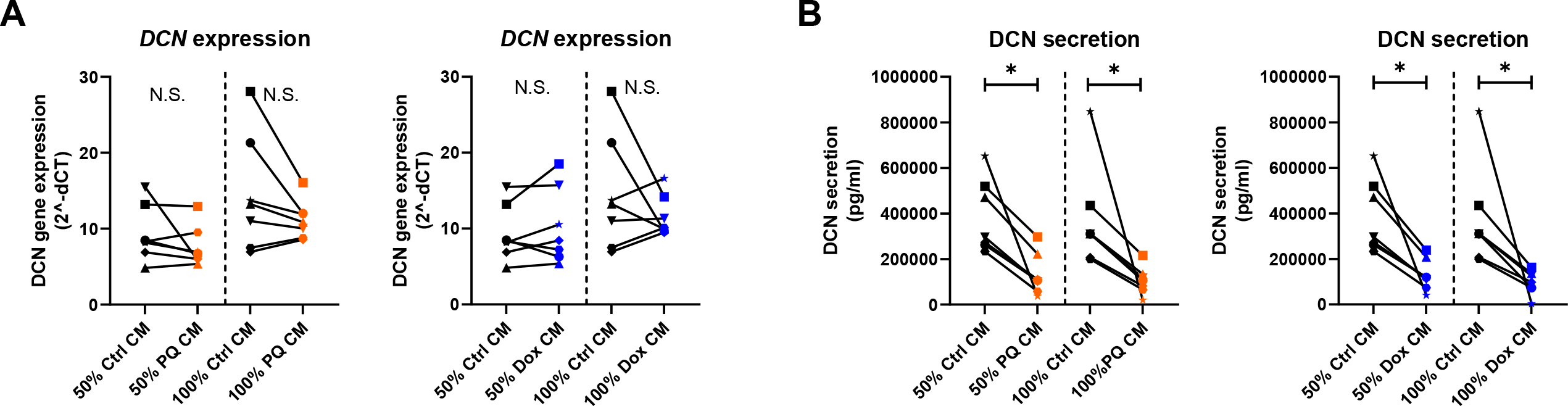
The effect of senescent CM on DCN secretion. Untreated SEO-COPD-derived fibroblasts were stimulated with control or senescent (PQ or Dox) conditioned medium (CM). *DCN* mRNA levels were measured after 24 hours **(A)**. DCN secretion was assessed in cell culture medium through ELISA after four days. DCN levels in CM itself (Supl. Fig. S5C) were subtracted from measured DCN levels **(B)**. Wilcoxon matched pairs signed-rank test was applied for statistical testing. * p<0.05.

*DCN* gene expression remained unchanged after 24 hours of t-PA stimulation (Figure 4A), whereas similar to CM stimulation, secreted DCN levels were significantly reduced after four days of 0.5nM and 10nM t-PA stimulation (Figure 4B).

**Figure 4.**
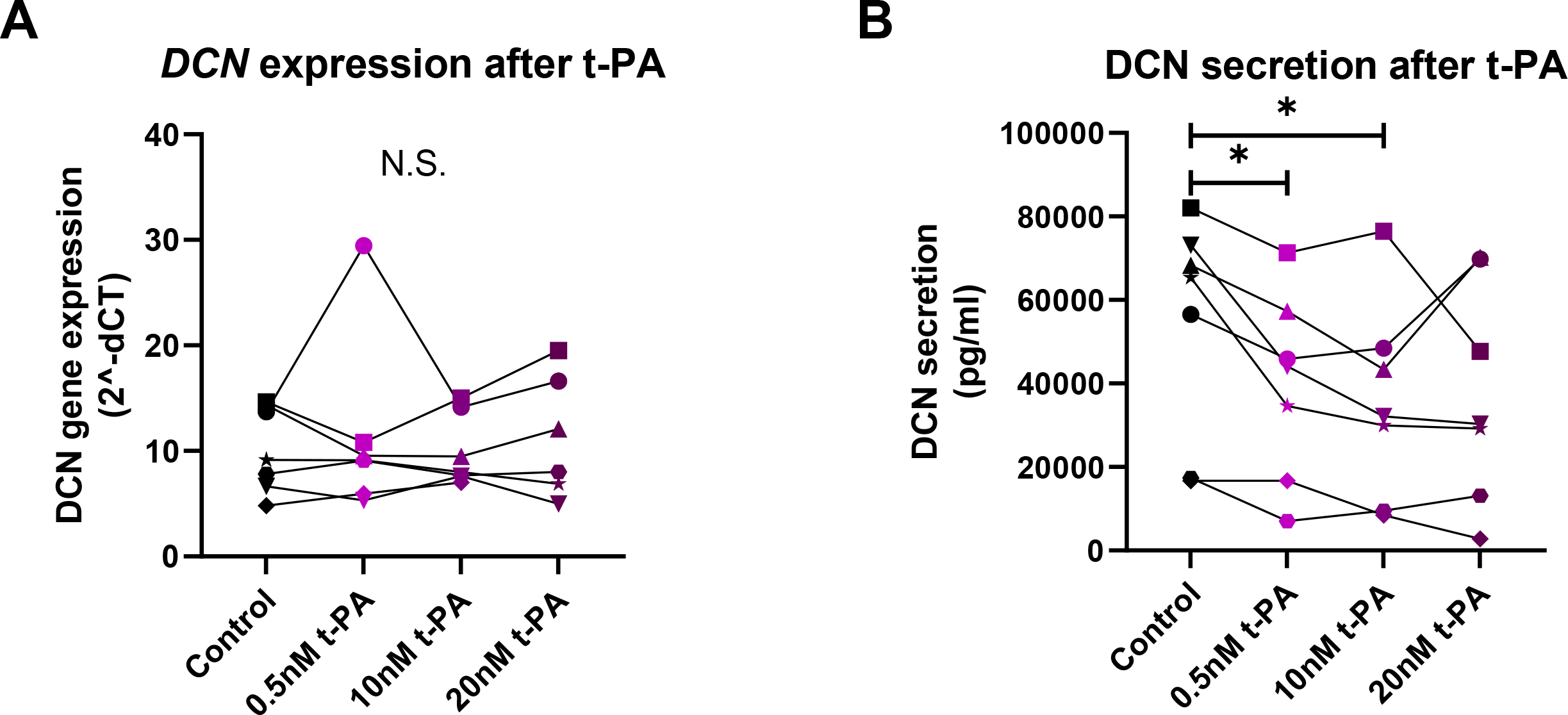
The effect of CM component t-PA on DCN secretion. Untreated SEO-COPD-derived fibroblasts were stimulated with 0.5nM, 10nM or 20nM tissue Plasminogen Activator (t-PA). *DCN* gene expression was measured after 24 hours **(A)**. After four days, DCN secretion was assessed in cell culture medium through ELISA **(B)**. Wilcoxon matched pairs signed-rank test was applied for statistical testing. * p<0.05.

## Discussion

Our study demonstrated that the SASP from senescence-induced SEO-COPD-derived fibroblasts induces a strong pro-inflammatory response and reduces DCN secretion in fibroblasts from the same patients. Treatment with the COPD-associated SASP protein t-PA also showed decreased DCN secretion, suggesting that this effect of CM is, at least partly, induced by t-PA. These findings imply a role for the SASP from senescent lung fibroblasts in chronic inflammation and tissue remodeling, two core features of COPD pathology. Importantly, our study was performed in lung fibroblasts derived from SEO-COPD patients, who developed a severe form of COPD at a relatively young age. The observed effects of senescence-induced SEO-COPD-derived fibroblasts, and their SASP, are in line with the theory that SEO-COPD patients undergo accelerated lung ageing and indicate that cellular senescence could be one of the mechanisms underlying SEO-COPD.

Our results demonstrate that the SASP from senescence-induced lung fibroblasts exerts pro-inflammatory effects on SEO-COPD-derived fibroblasts by increased IL-6 and IL-8 secretion. Previous research in a breast cancer cell line has shown that senescent CM stimulation increased IL-6 and IL-8 production, and that this increase was abolished in the presence of IL-6 or IL-8 antibodies, suggesting a positive feedback loop for IL-6 and IL-8 production(24). Our results in lung fibroblasts are consistent with this, as we observed increased levels of both cytokines in senescent CM (Supplementary Figure S5A & B), and in cell culture medium of senescent CM stimulated fibroblasts. To our knowledge, this study is the first to show that SASP from senescent fibroblasts induces a pro-inflammatory response in COPD. IL-6 and IL-8 are known to be elevated in COPD patients and have been linked to airway inflammation, exacerbations, and lung function decline(25–27). Hence, this potential positive feedback loop caused by senescent cells could contribute to the persistent inflammation observed in COPD lungs and could (partly) explain why COPD continues to progress after smoking cessation.

We observed reduced secreted DCN levels after senescent CM and t-PA stimulation, while *DCN* gene expression remained unaffected. Hence, the SASP, partly mediated by t-PA, could contribute to DCN protein degradation. DCN is a proteoglycan known to influence ECM properties by its role in collagen cross-linking(28) and regulation of locally active Transforming Growth Factor Beta(29, 30). DCN can be cleaved by MMP-2, MMP-3, MMP-7, and MMP-14(31, 32). T-PA can induce MMP-2 and MMP-3(17, 18). Hence, we speculate that the lowered DCN levels are caused by protein degradation via MMPs, either through direct or indirect effects of the SASP, as proteases are also known to be SASP components (13). Further studies are required to confirm the role of t-PA in DCN degradation. Our results are in line with previous findings on DCN in COPD, as lower DCN levels have been shown in (severe) emphysematous lungs(10). Altogether these findings suggest that senescent fibroblasts, and their SASP, reduce DCN levels, thereby playing an important role in ECM dysregulation in COPD.

In our experiments, senescent CM showed differential effects on paracrine senescence with increased percentage of SA-β-gal positive fibroblasts with both models of senescence-induced CM, and decreased *LMNB1* gene expression(33, 34), indicative of elevated cellular senescence, while the observed decreased p21 and p16 gene expression indicate reduced senescence(35) and cell numbers remained unaffected. Thereby, the effects on gene expression level were only observed after stimulation with PQ-induced senescent CM, suggesting that these effects might be dependent on the senescence inducer. As t-PA stimulation did not affect any of the senescence markers, t-PA is not likely to contribute to the observed alterations. SASP from COPD-derived fibroblasts comprises a mixture of, among others, inflammatory factors, and growth factors, which can have similar, synergistic, and opposing effects(14). When receiving CM, each fibroblast is exposed to a particular combination of SASP factors, causing a heterogeneous response.

We recognize that studying the effect of SASP through a CM model does not fully reflect the continuous exposure to SASP molecules that lung fibroblasts undergo *in vivo*. However, by using primary fibroblasts and two senescence induction models, we tried to model the *in vivo* situation as close as we can in a 2D *in vitro* model. Importantly, we showed that the effects on inflammation and ECM regulation were consistent with both senescence induction models, PQ and Dox, which induce cellular senescence via different mechanisms (oxidative stress(36, 37) and DNA damage(38, 39) respectively), that are hypothesized to be drivers of senescence in COPD-derived lung fibroblasts(12). Therefore, it is likely that the observed effects are due to increased numbers of senescent cells producing SASP and not due to compound specific properties.

Altogether, this study demonstrated that the SASP from senescent lung fibroblasts can promote inflammation and ECM remodeling in (SEO-)COPD. The obtained findings invite for future studies investigating the effects of senescent fibroblast-derived SASP, and individual SASP proteins, on other cell types relevant in COPD pathology through using advanced cell culture systems like co-culture models and 3D organoid cultures. Better understanding of the role of senescent lung fibroblasts and their SASP in COPD lungs could aid in identifying novel effective therapeutic targets. Specifically targeting (the negative effects) of senescent fibroblasts, for example by the promising senolytic therapies, might slow down accelerated lung ageing and consequently disease progression in COPD lungs.

## Acknowledgements

We would like to thank Marjan Reinders-Luinge (Department of Pathology and Medical Biology, University of Groningen, University Medical Center Groningen) for isolating primary parenchymal lung fibroblasts from patient-derived lung tissues.

## Author contributions

C.-A.B., W.T., M.v.d.B. & R.R.W. conceived and designed research; R.A.M., W.K. & R.R.W. performed experiments; R.A.M. & R.R.W. analyzed data; R.A.M., C.-A.B., W.T., M.v.d.B. & R.R.W. interpreted results of experiments; R.A.M. prepared figures; R.A.M. & R.R.W. drafted manuscript; R.A.M., C.-A.B., W.K., W.T., M.v.d.B. & R.R.W. edited and revised manuscript and approved final version of manuscript.

## References

1. W.H.O. The top 10 causes of death [Online]. 2019. https://www.who.int/news-room/fact-sheets/detail/the-top-10-causes-of-death [24 Mar. 2023].

2. Lundbäck B, Lindberg A, Lindström M, Rönmark E, Jonsson AC, Jönsson E, Larsson L-G, Andersson S, Sandström T, Larsson K. Not 15 But 50% of smokers develop COPD? - Report from the Obstructive Lung Disease in Northern Sweden Studies. Respir Med 97: 115–122, 2003. doi: 10.1053/rmed.2003.1446.

3. Baur X, Bakehe P, Vellguth H. Bronchial asthma and COPD due to irritants in the workplace - An evidence based approach. Journal of Occupational Medicine and Toxicology 7: 2012.

4. Rabe KF, Watz H. Chronic obstructive pulmonary disease. Lancet 389: 1931–1940, 2017. doi: 10.1016/S0140-6736(17)31222-9.

5. Brandsma C-A, De Vries M, Costa R, Woldhuis RR, Königshoff M, Timens W. Lung ageing and COPD: is there a role for ageing in abnormal tissue repair? Eur Respir Rev 26: 170073, 2017. doi: 10.1183/16000617.0073.

6. Silverman EK, Chapman HA, Drazen JM, Weiss ST, Rosner B, Campbell EJ, O’Donnell WJ, Reilly JJ, Ginns L, Mentzer S, Wain J, Speizer FE. Genetic epidemiology of severe, Early-onset Chronic Obstructive Pulmonary Disease. Am J Respir Crit Care Med 157: 1770–1778, 1998. doi: 10.1164/ajrccm.157.6.9706014.

7. Meiners S, Eickelberg O, Königshoff M. Hallmarks of the ageing lung. European Respiratory Journal 45: 807–827, 2015. doi: 10.1183/09031936.00186914.

8. Black PN, Ching PST, Beaumont B, Ranasinghe S, Taylor G, Merrilees MJ. Changes in elastic fibres in the small airways and alveoli in COPD. European Respiratory Journal 31: 998–1004, 2008. doi: 10.1183/09031936.00017207.

9. Segura-Valdez L, Pardo A, Gaxiola M, Uhal BD, Becerril C, Selman M. Upregulation of Gelatinases A and B, Collagenases 1 and 2, and Increased Parenchymal Cell Death in COPD. Chest 117: 684–694, 2000. doi: 10.1378/chest.117.3.684.

10. van Straaten JF, Coers W, Noordhoek JA, Huitema S, Flipsen JT, Kauffman HF, Timens W, Postma DS. Proteoglycan changes in the extracellular matrix of lung tissue from patients with pulmonary emphysema. Mod Pathol 12: 697–705, 1999.

11. Gey C, Seeger K. Metabolic changes during cellular senescence investigated by proton NMR-spectroscopy. Mech Ageing Dev 134: 130–138, 2013. doi: 10.1016/j.mad.2013.02.002.

12. Woldhuis RR, De Vries M, Timens W, Van Den Berge M, Demaria M, Oliver BGG, Heijink IH, Brandsma C-A. Link between increased cellular senescence and extracellular matrix changes in COPD. Am J Physiol Lung Cell Mol Physiol 319, 2020. doi: 10.1152/ajplung.00028.2020.-Chronic.

13. Coppé JP, Desprez PY, Krtolica A, Campisi J. The senescence-associated secretory phenotype: The dark side of tumor suppression. Annual Review of Pathology: Mechanisms of Disease 5: 99–118, 2010. doi: 10.1146/annurev-pathol-121808-102144.

14. 1. Woldhuis RR, Heijink IH, Van Den Berge M, Timens W, Oliver BGG, De Vries M, Brandsma CA. COPD-derived fibroblasts secrete higher levels of senescence-associated secretory phenotype proteins. Thorax 76: 508–511, 2021. doi: 10.1136/thoraxjnl-2020-215114.

15. Mori K, Dwek RA, Downing AK, Opdenakker G, Rudd PM. The activation of type 1 and type 2 plasminogen by type I and type II tissue plasminogen activator. Journal of Biological Chemistry 270: 3261–3267, 1995. doi: 10.1074/jbc.270.7.3261.

16. Yenari MA, Palmer JT, Bracci PM, Steinberg GK. Thrombolysis with tissue plasminogen activator is temperature dependent. Thromb Res 77: 475–481, 1995. doi: 10.1016/0049-3848(95)93883-2.

17. Song H, Cheng Y, Bi G, Zhu Y, Jun W, Ma W, Wu H. Release of matrix metalloproteinases-2 and 9 by S-nitrosylated caveolin-1 contributes to degradation of extracellular matrix in tPA-treated hypoxic endothelial cells. PLoS One 11, 2016. doi: 10.1371/journal.pone.0149269.

18. Suzuki Y, Nagai N, Yamakawa K, Kawakami J, Lijnen HR, Umemura K. Tissue-type plasminogen activator (t-PA) induces stromelysin-1 (MMP-3) in endothelial cells through activation of lipoprotein receptor-related protein. Blood 114: 3352–3358, 2009. doi: 10.1182/blood-2009-02-203919.

19. Noordhoek JA, Postma DS, Chong LL, Menkema L, Kauffman HF, Timens W, Van Straaten JFM, Van Der Geld YM. Different modulation of decorin production by lung fibroblasts from patients with mild and severe emphysema. COPD: Journal of Chronic Obstructive Pulmonary Disease 2: 17–25, 2005. doi: 10.1081/COPD-200050678.

20. Chen L, Hou J, Fu X, Chen X, Wu J, Han X. tPA promotes the proliferation of lung fibroblasts and activates the Wnt/β-catenin signaling pathway in idiopathic pulmonary fibrosis. Cell Cycle 18: 3137–3146, 2019. doi: 10.1080/15384101.2019.1669997.

21. Hao S, Shen H, Hou Y, Mars WM, Liu Y. tPA is a potent mitogen for renal interstitial fibroblasts: Role of β1 integrin/focal adhesion kinase signaling. American Journal of Pathology 177: 1164–1175, 2010. doi: 10.2353/ajpath.2010.091269.

22. Hu K, Yang J, Tanaka S, Gonias SL, Mars WM, Liu Y. Tissue-type plasminogen activator acts as a cytokine that triggers intracellular signal transduction and induces matrix metalloproteinase-9 gene expression. Journal of Biological Chemistry 281: 2120–2127, 2006. doi: 10.1074/jbc.M504988200.

23. Muro AF, Moretti FA, Moore BB, Yan M, Atrasz RG, Wilke CA, Flaherty KR, Martinez FJ, Tsui JL, Sheppard D, Baralle FE, Toews GB, White ES. An essential role for fibronectin extra type III domain A in pulmonary fibrosis. Am J Respir Crit Care Med 177: 638–645, 2008. doi: 10.1164/rccm.200708-1291OC.

24. Ortiz-Montero P, Londoño-Vallejo A, Vernot JP. Senescence-associated IL-6 and IL-8 cytokines induce a self- and cross-reinforced senescence/inflammatory milieu strengthening tumorigenic capabilities in the MCF-7 breast cancer cell line. Cell Communication and Signaling 15, 2017. doi: 10.1186/s12964-017-0172-3.

25. Donaldson GC, Seemungal TAR, Patel IS, Bhowmik A, Wilkinson TMA, Hurst JR, Maccallum PK, Wedzicha JA. Airway and Systemic Inflammation and Decline in Lung Function in Patients With COPD. Chest 128: 1995–2004, 2005. doi: 10.1378/chest.128.4.1995.

26. 1. Pesci A, Balbi B, Majori M, Cacciani G, Bertacco S, Alciato P, Donner CF. Inflammatory cells and mediators in bronchial lavage of patients with chronic obstructive pulmonary disease. European Respiratory Journal 12: 380–386, 1998. doi: 10.1183/09031936.98.12020380.

27. Aaron SD, Angel JB, Lunau M, Wright K, Fex C, Le Saux N, Dales RE. Granulocyte Inflammatory Markers and Airway Infection during Acute Exacerbation of Chronic Obstructive Pulmonary Disease. Am J Respir Crit Care Med 163: 349–355, 2001. doi: 10.1164/ajrccm.163.2.2003122.

28. Danielson KG, Baribault H, Holmes DF, Graham H, Kadler KE, Iozzo R V. Targeted Disruption of Decorin Leads to Abnormal Collagen Fibril Morphology and Skin Fragility. J Cell Biol 136: 729–743, 1997.

29. Markmann A, Hausser H, Schonherr E, Kresse H. Influence of decorin expression on transforming growth factor-mediated collagen gel retraction and biglycan induction. Matrix Biology 19: 631636, 2000.

30. Ferdous Z, Wei VM, Iozzo R, Höök M, Grande-Allen KJ. Decorin-transforming growth factor-β interaction regulates matrix organization and mechanical characteristics of three-dimensional collagen matrices. Journal of Biological Chemistry 282: 35887–35898, 2007. doi: 10.1074/jbc.M705180200.

31. Imai K, Hiramatsu A, Fukushima D, Pierschbacher MD, Okada Y. Degradation of decorin by matrix metalloproteinases[]: identification of the cleavage sites, kinetic analyses and transforming growth factor-β1 release. Biochem J 322: 809–814, 1997.

32. Mimura T, Han KY, Onguchi T, Chang JH, Kim TI, Kojima T, Zhou Z, Azar DT. MT1-MMP-Mediated cleavage of decorin in corneal angiogenesis. J Vasc Res 46: 541–550, 2009. doi: 10.1159/000226222.

33. Dimri GP, Lee X, Basile G, Acosta M, Scott G, Roskelley C, Medrano EE, Linskens M, Rubel I, Pereira-Smith O, Peacocke M, Campisi J. A biomarker that identifies senescent human cells in culture and in aging skin in vivo. Cell Biology 92: 9363–9367, 1995. doi: 10.1073/pnas.92.20.9363.

34. Freund A, Laberge RM, Demaria M, Campisi J. Lamin B1 loss is a senescence-associated biomarker. Mol Biol Cell 23: 2066–2075, 2012. doi: 10.1091/mbc.E11-10-0884.

35. Stein GH, Drullinger LF, Soulard A, Dulić V. Differential Roles for Cyclin-Dependent Kinase Inhibitors p21 and p16 in the Mechanisms of Senescence and Differentiation in Human Fibroblasts. Mol Cell Biol 19: 2109–2117, 1999. doi: 10.1128/mcb.19.3.2109.

36. Chinta SJ, Woods G, Demaria M, Rane A, Zou Y, McQuade A, Rajagopalan S, Limbad C, Madden DT, Campisi J, Andersen JK. Cellular Senescence Is Induced by the Environmental Neurotoxin Paraquat and Contributes to Neuropathology Linked to Parkinson’s Disease. Cell Rep 22: 930–940, 2018. doi: 10.1016/j.celrep.2017.12.092.

37. Bus JS, Gibson JE. Paraquat: Model for Oxidant-Initiated Toxicity. Environ Health Perspect 55: 37–46, 1984. doi: 10.1289/ehp.845537.

38. Hu X, Zhang H. Doxorubicin-induced cancer cell senescence shows a time delay effect and is inhibited by epithelial-mesenchymal transition (EMT). Medical Science Monitor 25: 3617–3623, 2019. doi: 10.12659/MSM.914295.

39. 1. Bilardi RA, Kimura KI, Phillips DR, Cutts SM. Processing of anthracycline-DNA adducts via DNA replication and interstrand crosslink repair pathways. Biochem Pharmacol 83: 1241–1250, 2012. doi: 10.1016/j.bcp.2012.01.029.

