## Supplementary material for "Strong paracrine effects of SASP from senescence-induced severe early-onset COPD-derived fibroblasts": Online Supplement

#### Subjects & ethical approval

Primary lung fibroblasts derived from explant material of seven SEO-COPD patients were collected in agreement with the Research Code of the University Medical Centre Groningen and national ethical and professional guidelines. The fibroblasts used in this study were derived from leftover lung material after lung surgery from archival materials that are exempt from consent with compliance with applicable laws and regulations. This material was not subject to the Medical Research Human Subjects Act in the Netherlands, and, therefore, an ethics waiver was provided by the Medical Ethical Committee of the University Medical Center Groningen. All samples and clinical information were pseudonymized before experiments were performed, blinding all directly identifying information to the investigators.

#### Gene expression

RNA was collected using TRIzol (Invitrogen, Carlsbad, US), isolated using the manufacturer's protocol and stored at -20°C until use. RNA concentrations were quantified via the Nanodrop ND-1000 Spectrophotometer (ThermoFisher Scientific). 200ng RNA per sample was used for cDNA synthesis using SuperScript II (Invitrogen) according to manufacturer's protocol(1). Quantitative Real-Time Polymerase Chain Reaction (qRT-PCR) was performed as described before to measure gene expression of *POLR2A*, *CDKN1A*, *CDKN2A*, *LMNB1*, *IL6*, *CXCL8*, *FN1*, *MMP2* and *DCN*(1). TaqMan assay IDs used for qRT-PCR measurements are shown in Supplementary Table 2. Samples were run in triplicates and 2-ΔCP values were calculated for individual genes in relation to the housekeeping gene *POL2RA*. Final data were visualized using GraphPad Prism 8 Software (Prism, Redmond, US).

#### Protein secretion

During CM collection, and four days after CM and t-PA stimulation, cell-free supernatants were collected to measure levels of secreted Interleukin (IL-)6, IL-8 and DCN via Enzyme-Linked Immunosorbent Assays (ELISAs). Supernatants were stored at -80°C until use. ELISAs were performed using manufacturer's protocol (R&D Systems, Minneapolis, US). Obtained data were interpolated and visualized using GraphPad Prism 8 Software. DCN levels in CM itself (Supplementary figure 3C) were subtracted from measured DCN levels in supernatants after CM stimulation to calculate final DCN levels.

#### Senescence-associated beta-galactosidase staining

Fibroblasts used for CM collection and fibroblasts four days after CM stimulation were stained for SA-β-gal activity to assess cellular senescence. Fibroblasts were fixed with 2% formaldehyde + 0.2% glutaraldehyde in PBS for five minutes, and stained as described previously(2). Hematoxylin was used to counterstain nuclei. From each well, images were captured at four random locations at a final magnification of 100X using a TissueFAXS microscope (TissueGnostics GmbH, Vienna, Austria). The number of nuclei and SA-β-gal positive cells were counted using ImageJ to calculate the average percentage of SA-β-gal positive cells per well.

**Supplementary table 1. Patient characteristics**

| Variable | n | Age | Male / Female (n) | Pack-years | Stop months | $FEV_1$ % predicted | FVC % predicted | $FEV_1$ / FVC |
|---|---|---|---|---|---|---|---|---|
| **Median (Range)** | 7 | 51 (48 - 53) | 3/4 | 30 (8 - 42) | 84 (16 – 96) | 15 (12 - 28) | 41 (25 - 78) | 26 (23 - 45) |

Data are shown as median and range. $FEV_1$, Forced Expiratory Volume in one second; FVC, Forced Vital Capacity.

**Supplementary table 2. TaqMan assays used for gene expression measurements**

| Gene name | Protein | Company | Assay ID |
|---|---|---|---|
| *POLR2A* | RNA polymerase II subunit A | ThermoFisher Scientific | Hs00172187_m1 |
| *CDKN1A* | P21 | ThermoFisher Scientific | Hs00355782_m1 |
| *CDKN2A* | P16 | ThermoFisher Scientific | Hs00923894_m1 |
| *LMNB1* | Lamin B1 | ThermoFisher Scientific | Hs01059210_m1 |
| *IL6* | Interleukin 6 | ThermoFisher Scientific | Hs00174131_m1 |
| *CXCL8* | Interleukin 8 | ThermoFisher Scientific | Hs00174103_m1 |
| *DCN* | Decorin | ThermoFisher Scientific | Hs00370385_m1 |
| *FN1* | Fibronectin 1 | ThermoFisher Scientific | Hs00365052_m1 |
| *MMP2* | Matrix Metalloproteinase-2 | ThermoFisher Scientific | Hs01548727_m1 |

For each TaqMan assay, the corresponding gene name, encoded protein, company, and Assay ID are presented.

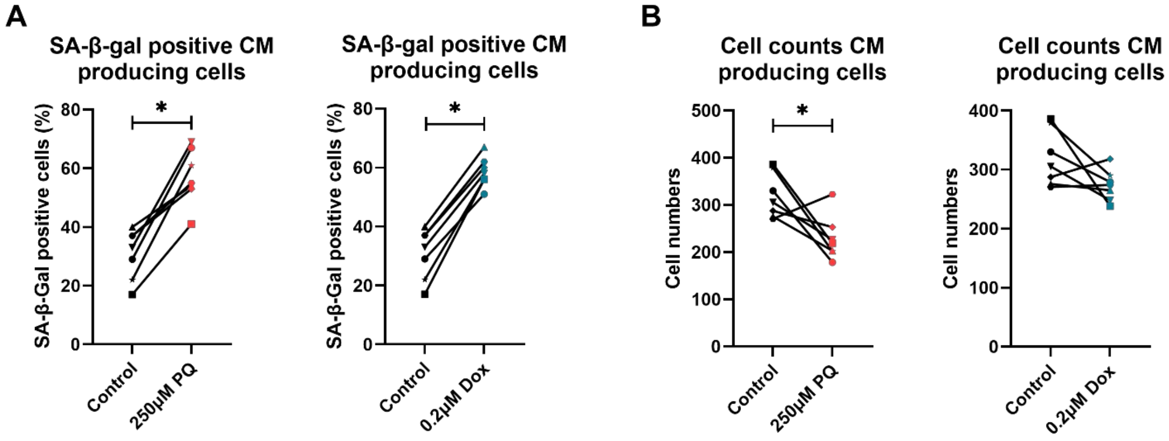


**Supplementary Figure 1. SA-β-gal staining scores of CM producing cells.** SEO-COPD-derived lung fibroblasts were treated with 250μM PQ or 0.2μM Dox or were kept as control and reseeded in 12-well plates for CM production. The percentage of SA-β-gal staining positive cells **(A)** and total number of cells **(B)** were assessed at the moment of CM collection, i.e. three days after reseeding the fibroblasts. Wilcoxon matched pairs signed rank test was applied for statistical testing. * $p < 0.05$. PQ, Paraquat; Dox, Doxorubicin; SA-β-gal, Senescence Associated Beta Galactosidase.

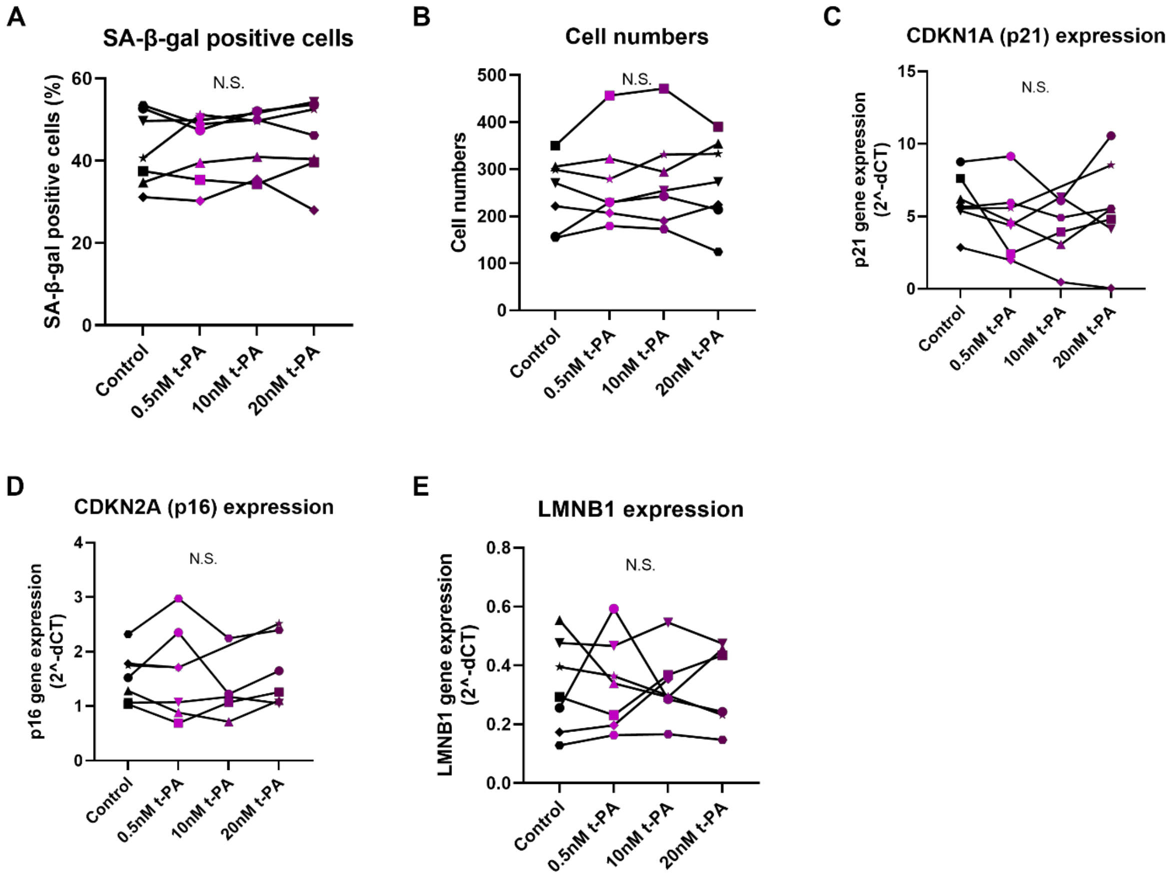


**Supplementary Figure 2. The effect of t-PA on paracrine senescence.** Untreated SEO-COPD-derived lung fibroblasts were stimulated with 0.5nM, 10nM or 20nM t-PA. The percentage of SA-β-gal staining positive cells **(A)** and total number of cells **(B)** were assessed after four days. p21 (CDKN1A) **(C)**, p16 (CDKN2A) **(D)** and *LMNB1* **(E)** mRNA expression was measured after 24 hours. Wilcoxon matched pairs signed rank test was applied for statistical testing. * $p < 0.05$. t-PA, tissue Plasminogen Activator; SA-β-gal, Senescence Associated Beta Galactosidase.

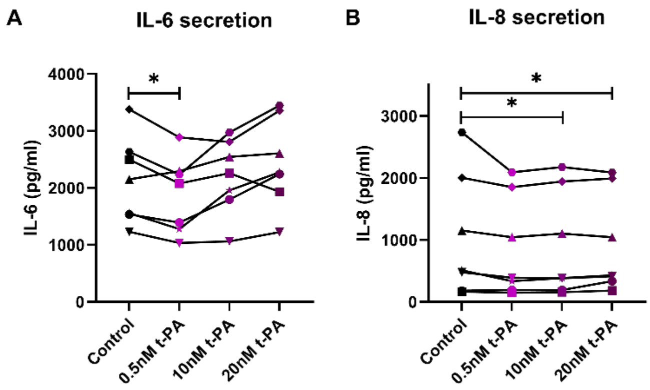


**Supplementary Figure 3. The effect of t-PA on pro-inflammatory interleukin secretion.** Untreated SEO-COPD-derived lung fibroblasts were stimulated with 0.5nM, 10nM or 20nM t-PA. After four days, IL-6 **(C)** and IL-8 **(D)** secretion was assessed in cell culture medium using ELISA. Wilcoxon matched pairs signed rank test was applied for statistical testing. * $p < 0.05$. t-PA, tissue Plasminogen Activator; IL-6, Interleukin 6; IL-8, Interleukin 8.

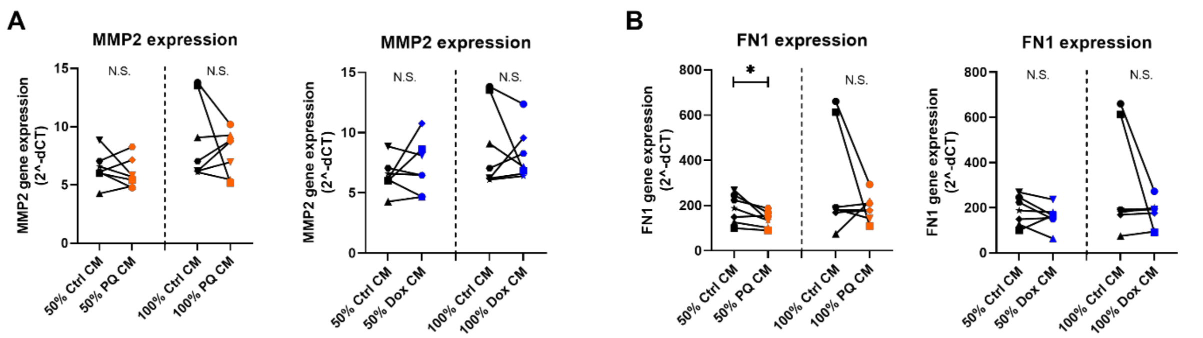


**Supplementary Figure 4. The effect of senescent CM on extracellular matrix gene expression.** SEO-COPD-derived lung fibroblasts were treated with control or (PQ- or Dox-induced) senescent CM. *MMP2* **(A)** and *FN1* **(B)** mRNA expression was measured by qRT-PCR after 24 hours of CM treatment. Wilcoxon matched pairs signed rank test was applied for statistical testing. * $p < 0.05$. PQ, Paraquat; Dox, Doxorubicin; MMP2, Matrix metalloproteinase-2; FN1, Fibronectin 1.

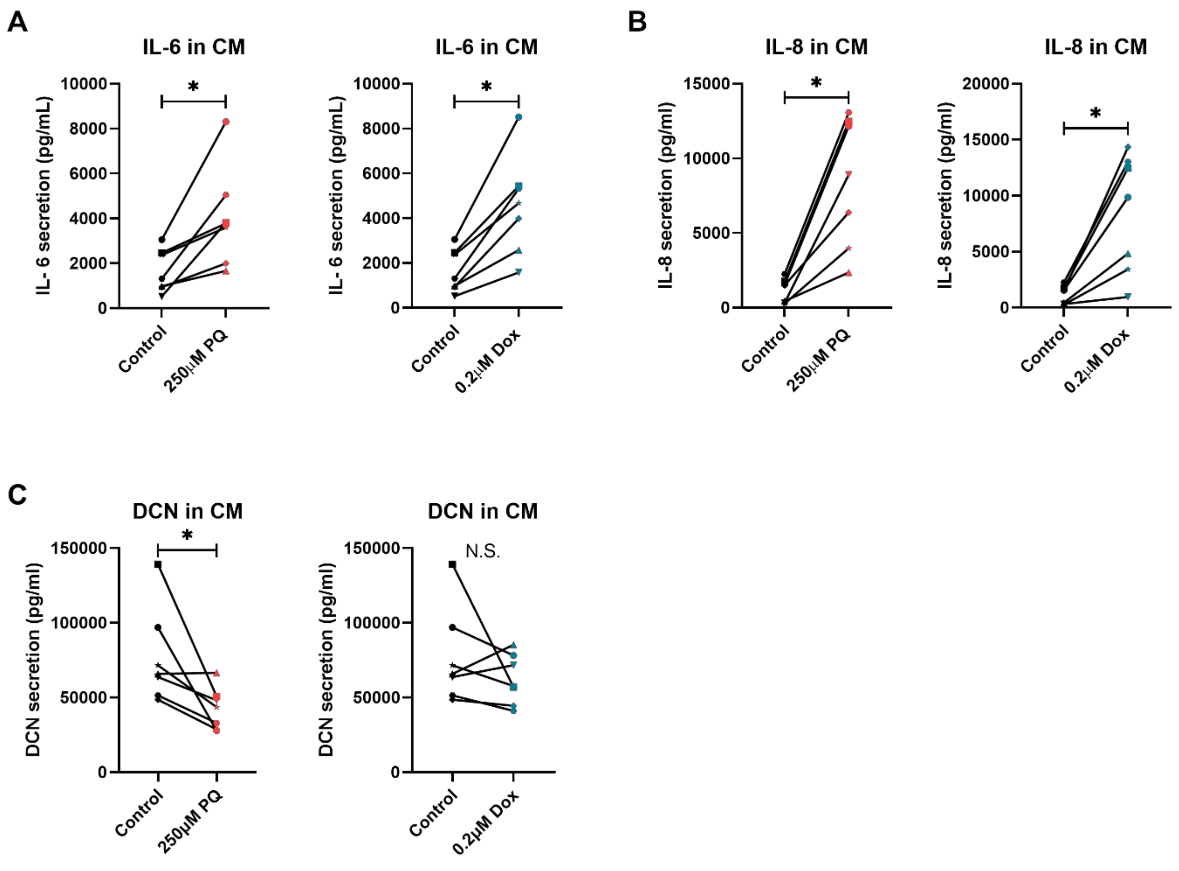


**Supplementary Figure 5. IL-6, IL-8 and DCN levels in CM.** SEO-COPD-derived lung fibroblasts were treated with 250µM PQ or 0.2µM Dox or were kept as control and reseeded in 12-well plates for CM production. Il-6 **(A),** IL-8 **(B)** and DCN secretion **(C)** were assessed in the collected CM. Wilcoxon matched pairs signed rank test was applied for statistical testing. * $p < 0.05$. PQ, Paraquat; Dox, Doxorubicin; IL-6, Interleukin 6; IL-8, Interleukin 8; DCN, Decorin.

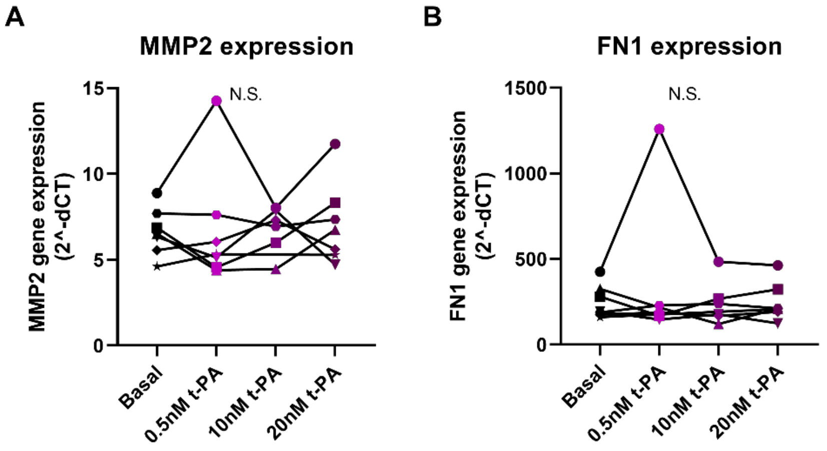


**Supplementary Figure 6. The effect of t-PA on extracellular matrix gene expression.** Untreated SEO-COPD-derived lung fibroblasts were stimulated with 0.5nM, 10nM or 20nM t-PA. After 24 hours, *MMP2* **(A)** and *FN1* **(B)** mRNA expression was measured by qRT-PCR. Wilcoxon matched pairs signed rank test was applied for statistical testing. * $p < 0.05$. t-PA, tissue Plasminogen Activator; MMP2, Matrix metalloproteinase-2; FN1, Fibronectin 1.

### References


1. **Woldhuis RR**, **De Vries M**, **Timens W**, **Van Den Berge M**, **Demaria M**, **Oliver BGG**, **Heijink IH**, **Brandsma C-A**. Link between increased cellular senescence and extracellular matrix changes in COPD. *Am J Physiol Lung Cell Mol Physiol* 319, 2020. doi: 10.1152/ajplung.00028.2020.-Chronic.

2. **Dimri GP**, **Lee X**, **Basile G**, **Acosta M**, **Scott G**, **Roskelley C**, **Medrano EE**, **Linskens M**, **Rubel I**, **Pereira-Smith O**, **Peacocke M**, **Campisi J**. A biomarker that identifies senescent human cells in culture and in aging skin in vivo. *Cell Biology* 92: 9363–9367, 1995. doi: 10.1073/pnas.92.20.9363.
